# Identification and Mechanistic Basis of non-ACE2 Blocking Neutralizing Antibodies from COVID-19 Patients with Deep RNA Sequencing and Molecular Dynamics Simulations

**DOI:** 10.1101/2022.06.29.498206

**Authors:** Alger M. Fredericks, Kyle W. East, Yuanjun Shi, Jinchan Liu, Federica Maschietto, Alfred Ayala, William G. Cioffi, Maya Cohen, William G. Fairbrother, Craig T. Lefort, Gerard J. Nau, Mitchell M. Levy, Jimin Wang, Victor S. Batista, George P. Lisi, Sean F. Monaghan

## Abstract

Variants of severe acute respiratory syndrome coronavirus-2 (SARS-CoV-2) continue to cause disease and impair the effectiveness of treatments. The therapeutic potential of convergent neutralizing antibodies (NAbs) from fully recovered patients has been explored in several early stages of novel drugs. Here, we identified initially elicited NAbs (Ig Heavy, Ig lambda, Ig kappa) in response to COVID-19 infection in patients admitted to the intensive care unit at a single center with deep RNA sequencing (>100 million reads) of peripheral blood as a diagnostic tool for predicting the severity of the disease and as a means to pinpoint specific compensatory NAb treatments. Clinical data were prospectively collected at multiple time points during ICU admission, and amino acid sequences for the NAb CDR3 segments were identified. Patients who survived severe COVID-19 had significantly more of a Class 3 antibody (C135) to SARS-CoV-2 compared to non-survivors (16,315 reads vs 1,412 reads, p=0.02). In addition to highlighting the utility of RNA sequencing in revealing unique NAb profiles in COVID-19 patients with different outcomes, we provided a physical basis for our findings via atomistic modeling combined with molecular dynamics simulations. We established the interactions of the Class 3 NAb C135 with the SARS-CoV-2 spike protein, proposing a mechanistic basis for inhibition via multiple conformations that can effectively prevent ACE2 from binding to the spike protein, despite C135 not directly blocking the ACE2 binding motif. Overall, we demonstrate that deep RNA sequencing combined with structural modeling offers the new potential to identify and understand novel therapeutic(s) NAbs in individuals lacking certain immune responses due to their poor endogenous production. Our results suggest a possible window of opportunity for administration of such NAbs when their full sequence becomes available. A method involving rapid deep RNA sequencing of patients infected with SARS-CoV-2 or its variants at the earliest infection time could help to develop personalized treatments using the identified specific NAbs.

## Introduction

Severe acute respiratory syndrome coronavirus 2 (SARS-CoV-2), responsible for coronavirus disease 2019 (COVID-19) continues to cause critical illness requiring intensive care unit (ICU) admission, which is currently driven, in part, by variants.^1^ Few direct treatments against COVID-19 are available^2^ and with variants comes further potential for escape from current treatments and vaccines.^3,4^ Variants have caused an increase in cases, led to recommendations for additional vaccine doses, and blunted the efficacy of two current antibody treatments. Quickly identifying novel antibodies to target the new variants could aid in the care of these patients, reducing viral load and preventing hypoxemia. This is important on two levels; first because the majority of antibodies target the SARS-CoV-2 spike protein receptor-binding domain (RBD) and mutations in this region are present in variants,^5,6^ and second, because the *de novo* discovery and testing of antibodies is typically a very time-consuming process.^7^

Long-lived, strongly neutralizing antibodies (Nabs) against the spike protein of SARS-CoV-2 have been developed for therapeutic use,^8-11^ and the number of studies of potential Nabs increases rapidly due to well-developed methodologies for single-cell RNA sequencing and production of monoclonal antibodies with predefined specificity.^12-14^ The measured binding affinity for some of these NAb-spike protein complexes is as tight as 7.2 *p*mol (*K*_D_),^11^ about three-orders of magnitude tighter than the ACE2-spike complex. Most of these NAbs bind the receptor-binding motif (RBM) site of the spike protein receptor-binding domain (RBD), *i*.*e*. the ACE2-binding site, and sterically block ACE2 binding. These “ACE2 blocking” NAbs can easily strip bound ACE2 from a spike protein complex to free the virus from host cell attachment.

The binding of multiple ACE2 blocking NAbs to three “up” positioned RBD opens the central pore of the spike trimer, permitting the central stalk of the S2 fragment of the spike protein to extend (*i*.*e*. the postfusion state) relative to the bent prefusion state.^15-17^ When this occurs away from a host cell membrane, it permanently disarms the spike trimer and also exposes more epitopes for binding of additional NAbs. However, the ACE2 blocking NAbs cannot bind the spike protein when its three RBDs are in “down” positions. Thus, immunity must also rely on NAbs of other types that bind different regions of the RBD or different domains of the spike protein entirely, which are often referred to as non-ACE2 blocking NAbs, a class of NAbs that have not been well studied.

In the treatment of viruses, NAb therapeutics are often cocktails of two noncompeting NAbs that can simultaneously bind, for example, different locations of the SARS-CoV-2 spike protein RBD, one at an ACE2 blocking site and the other at a non-ACE2 blocking site.^10,18-20^ The rationale for the latter NAb is to reduce the probability of spontaneous mutations that become resistant to ACE2 blocking. Because the ACE2 binding site is often conserved in variants of concern, most known spike mutations are located distal to this site and do not alter the binding affinity of ACE2-blocking NAbs significantly. Thus, it is presently unclear how non-ACE2 blocking NAbs work and whether they play a dominant role in NAb cocktails.

Efforts to identify both types of convalescent NAbs have been carried out using single-cell RNA sequencing (RNAseq) of B- or T-cell samples from recovered patients infected with SARS-CoV-2. At least one study included patients who did not survive.^21^ However, the systematic time-course evolution of NAbs in patients has not been evaluated previously, which became our motivation for this study. We used RNAseq data from critically ill COVID-19 patients in the ICU to identify novel antibodies elicited during the course of the disease, with particular focus on NAbs generated by patients who survived, to inform the severity of the disease and future antibody or vaccine development. We found that time-course evolution of NAbs differs between surviving and non-surviving COVID-19 patients. COVID-19 survivors expressed a very high level of non-ACE2 blocking NAbs at the time of initial infection that helped them completely fend off additional persistent infections. The level of non-ACE2 blocking NAbs was much higher than non-survivors, suggesting that non-ACE2 blocking NAbs, and not necessarily the convergent ACE2 blocking NAbs, are critical for recovery. We also used the recently proposed structural classification of COVID-19 antibodies^22^ to categorize potential efficacious antibodies and provide a possible rationale for this effect. Using NAb C135 as a representative of the non-ACE2 blocking class,^22,23^ we carried out full atomistic modeling and molecular dynamics (MD) simulations of the NAb-spike protein complex, the results of which provide a biophysical basis for how non-ACE2 blocking NAbs neutralize SARS-CoV-2.

## Materials and Methods

### Patient study design, population, and setting

ICU participants at a single tertiary care hospital were enrolled after themselves or surrogates provided informed consent (IRB Approval # 411616). SARS-CoV-2 infection was based on positive PCR from the nasopharynx. Patients were followed until discharge or death and clinical information was collected prospectively. Blood samples were collected on Day 0 of ICU admission and Day 3 whenever possible.

### RNA extraction, sequencing, data protection and quality assurance

Blood was collected directly from the patient into PAXgene tubes (Qiagen, Germantown, MD) on both Day 0 and Day 3 and was sent to Genewiz (South Plainfield, NJ) for RNA extraction, ribosomal RNA depletion, and sequencing on Illumina HiSeq machines with greater than 100 million reads per sample. In order to ensure security of the genomic data to HIPAA standards, the raw files were returned on password protected external hard drives and all analysis was done on servers within the hospital firewalls. As previously described with some of the data,^24^ quality was assessed by using FastQC.^25^ Sequencing data was aligned using STAR aligner with standard parameters. Unmapped reads were included in the output.

### Identification of antibodies

Alignment files were then parsed for reads mapping onto the V(D)J locus using ImReP^26^ to identify novel antibodies produced by the patients with active COVID-19 disease. The resulting CDR3 sequences were then compared across survivors and non-survivors retrospectively. Only sequences that appeared in every patient in each group were considered distinct to that group. Comparison was done using NCBI blast, with a threshold of 66% length match and 70% sequence match. Querying sequences by time point, across time points, further filtered blast output and survivors vs. non-survivors by time point and across time points.

### Classification of CDR3 regions and model of human neutralizing antibodies (NAb)

The patient-derived light chain CDR3 sequences were aligned to known CDR3 regions of previously identified neutralizing antibodies (**Supplemental Table 1**). CDR3 sequences were classified according to primary sequence similarity to the CDR3 classifications of known three-dimensional structures based on their interactions within NAb-spike protein complexes.^22^ If the primary sequence did not align to any of those sequences, the CDR3 was left unclassified.

### Statistical analysis

SigmaPlot 14.5 (Sysstat Software) was used for analysis. T-tests were used to compare survivors and non-survivors, but paired t-tests were used to compare across time points. Alpha was set at 0.05.

### Molecular dynamics simulations

A complete NAb-spike protein model was built for MD simulations primarily based on the cryo-EM map (emd22736) reported at 3.5-Å resolution, starting with the corresponding PDB 7k8z coordinates (**Figs. 1-4**) Any missing residues, particularly charged residues often located on surface loops, could incorrectly modulate MD trajectories. Rebuilding was therefore extensive due to the highly incomplete nature of the deposited 7k8z coordinates, which likely resulted from a lack of confidence in the experimental map at relatively low local resolution, particularly near the RBD/NAb C135 interface. Our rebuilt model included two missing loops (S443-G447 and E471-F489), one single-residue gap (V502), and many truncated large residues to those containing only a single Cβ sidechain atom (*i*.*e*. to alanine) even in the most ordered version of the RBDs, including R346 and N440 (these two sidechains were not present in the 7k8z coordinates although they were included in published figures). The first missing loop and the single-residue gap are located at the RBD/NAb C135 interface and the second missing loop is at the RBD/RBD interface mediated by glycans. Rebuilding also included the entire Fab fragment of NAb C135 for the “up” positioned RBD and half of the Fab fragment for each of the two “down” positioned RBDs (**Figs, S1, S2**). This structure contains one “up” and two “down” RBDs. The only variable loop-containing half of the C135 Fab fragment that is in direct contact with the RBD was included in our MD simulations (**Fig. S3**). A slightly truncated version of the RBD (N334 to L516, a few residues shorter than the conventional definition after excluding two paired strands at its N-and C-termini) as well as the glycan attached to N343 was included for the RBD in the RBD-NAb complex in our simulations.

**Figure 1.**
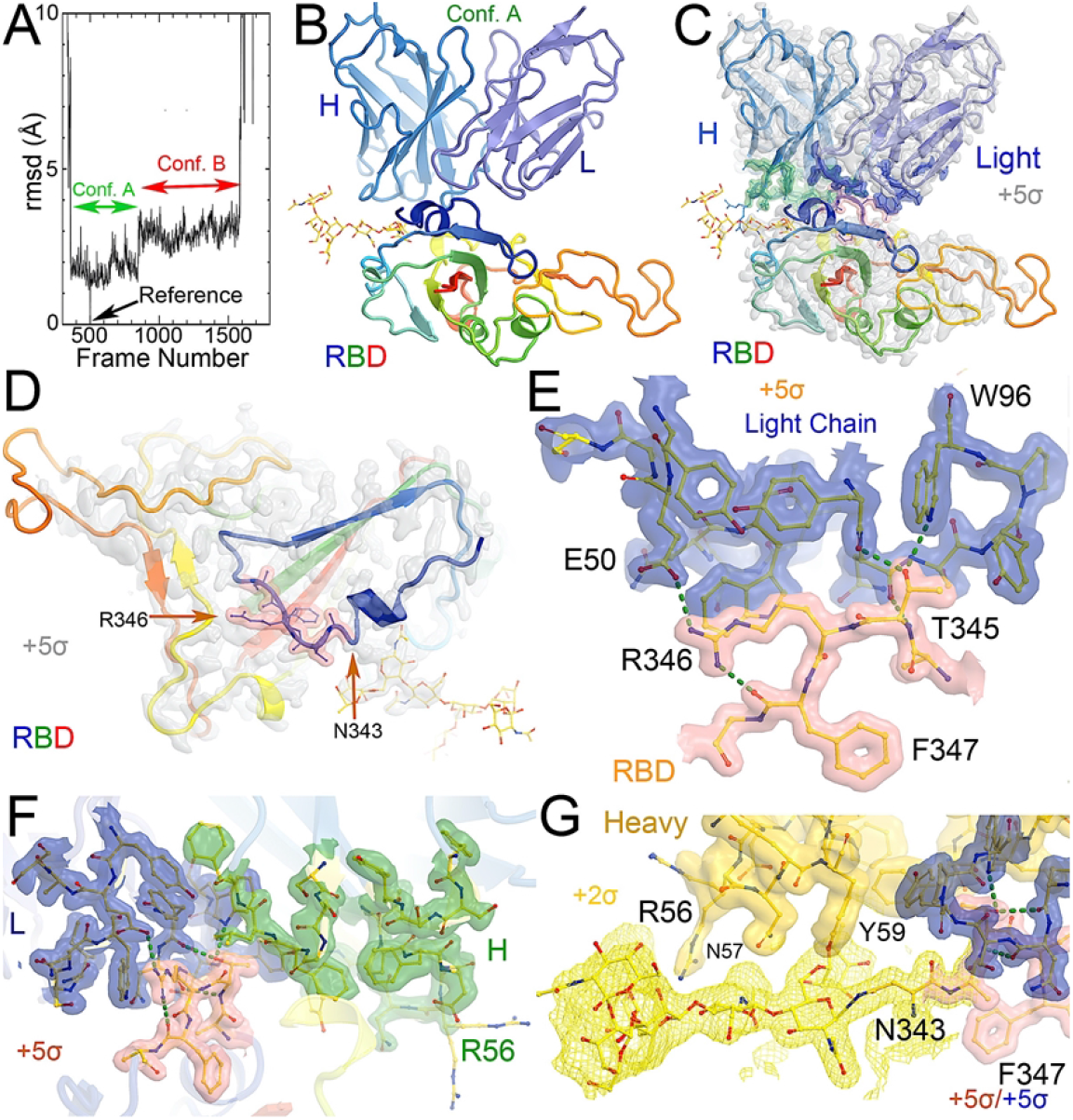
MD-derived electrostatic potential (ESP) maps and the equilibrated structure for NAb conformation A. (A) Frames corresponding to conformation A (green arrow) and conformation B (red arrow). (B) Overall structure of conformation A between the spike RBD (rainbow colors with glycosylated N343 in ball-and-stick) and heavy (slate)/light (light blue) chains of the C135 Fab. (C) MD-derived ESP maps contoured at +5σ. Contact interface maps are colored: light chain in blue, heavy chain in green, and RBD in pink. (D) Closeup view of the RBD with the map from (C). (E) Interactions between the RBD and the NAb light chain. (F) Interactions among RBD, heavy/light chains with maps contoured +5σ. (G). Reduced contouring level to +2.5σ for the heavy chain to show the features for N343 glycan.

The entire C135 Fab fragment bound to each of the three RBDs was clearly visible in the experimental map at +2σ contouring level for confident rebuilding (**Fig. S1, S2**). Although a high-resolution crystal structure of NAb C135 was docked into the cryo-EM map of the 7k8z complex, the conformation of this NAb C135 in the complex clearly differs in variable loop 3 of the heavy chain from that of the isolated Nab C135 structure, suggesting an induced-fit conformational change of this Fab fragment. This induced-fit conformational change was to justify additional MD simulations. Following complete RBD rebuilding, we mapped Omicron variant mutations and found that two of these are located at the RBD/NAb C135 interface and five additional mutations are nearby (**Fig. S4E, S4F**).

After obtaining the more complete coordinates, a glycan moiety was attached to N343 according to the experimental map using the Glycan Reader & Modeler module in CHARM-GUI.^27^ Topology and parameters were built generated using the PSFGEN tool in VMD.^28^ The system was solvated in the middle of TIP3P water box (∼ 104 Å X 104 Å X 104 Å). Five cycles of equilibration were run using NAMD.^29,30^ The MD simulations were run using a 2 fs time step and under 1.013225 bar (NPT ensemble). During the first cycle (500 ps), the entire system was frozen with the exception of waters and ions. For the second cycle (15 ns), the protein sidechains and residues within 8 Å of the C1325/RBD interface were released. For the third cycle (10ns), all restraints were released. For the fourth cycle (500 ps), the full released system was heated from 1 K to 298 K. The freed system from the equilibration run was taken as the starting point for the production run, which was carried out on GPU and used the AMBER force field parameters.^31^ ParmEd was used to convert the parameters and hydrogen mass repartitioning (HMR) was used to enable the MD simulations to employ 1 4 fs step.^31^ The overall production run was carried out for 465 ns. The resulting MD trajectories were analyzed using MD-derived ESP maps as described elsewhere.^32-34^

## Results

### A new pattern of time-course evolution of non-ACE2 blocking NAbs

A total of 15 patients had samples collected on Day 0 initially admitted in the ICU, 12 of which also had samples collected on Day 3 (2 patients were discharged from the ICU and one succumbed to COVID-19). Clinical and demographic data is previously reported^24^ and also presented in **Table 1**. We analyzed antibody (Ig) counts of Ig Heavy, Ig kappa, and Ig lambda chains in COVID-19 survivors and non-survivors (Table 1) in aggregate and compared across Days 0 and 3 time points. There was no difference in the number of Ig Heavy, Ig kappa, or Ig lambda counts between survivors and non-survivors on any given day, but when comparing *across time points*, we found a significant increase in Ig lambda across all patients (16,590 vs 33,610, p=0.009). When analyzing survivors alone, we observed the same trend (11,313 vs 31,087, p=0.0426), which was not present in non-survivors (21,868 vs 36,133, p=0.1504).

**Table 1.** Demographics and Antibody Types Demographic data and antibody types per group (survivor vs. non-survivor) at ICU Day 0 and ICU Day 3

| Table 1 Demographics and Antibody Types |  |  |  |  |
| --- | --- | --- | --- | --- |
|  | Day 0 |  | Day 3 |  |
|  | Survivor (8) | Non-Survivor (7) | Survivor (6) | Non-Survivor (6) |
| Median age (SD) | 53.9 (22.7) | 64.1(13.8) | 53.5 (18.5) | 60.9 (11.8) |
| Female - no. (%) | 2 (25) | 2 (29) | 2 (33) | 2 (33) |
| Race |  |  |  |  |
| White - no. (%) | 2 (25) | 4 (57) | 1 (17) | 3 (50) |
| Black or African American - no. (%) | 1 (13) | 1 (14) | 1 (17) | 1 (17) |
| Other Race - no. (%) | 5 (62) | 2 (29) | 4 (66) | 2 (33) |
| Ethnicity |  |  |  |  |
| Hispanic - no. (%) | 4 (50) | 5 (71) | 3 (50) | 4 (66) |
| Median BMI (SD) | 28.1 (4.1) | 31.5 (4.9) | 26.6 (3.6) | 31.7 (5.4) |
| Ig Heavy (SD) | 18,77.6 (31,136.3) | 17,247 (17,780) | 45,029.8 (48,919.6) | 39,982.1 (33,397.3) |
| Ig Kappa (SD) | 70,548.6 (136,600.2) | 43,630.7 (44,591.6) | 65,278.3 (83,406) | 34,784.3 (22,616.4) |
| Ig Lambda (SD) | 9,289.9 (11,302.4) | 19,000.9 (24,621.1) | 31,087.3 (29,572.1) | 36,133 (30,801) |
| Total Reads (SD) | 99,478.5 (178,615) | 80,942.6 (86,484.6) | 142,306.8 (141,072.9) | 111,939.2 (84,022.7) |

We analyzed the complementarity-determining region (CDR3) sequence counts of each patient (*i*.*e*. a variable amino acid sequence that NAbs use to recognize their antigen) and categorized the antibody types using a recently published classification system^22^ on Day 0 and Day 3 (Supplemental Tables 2 and 3, respectively). The resulting categorizations of antibody types, antibody classes, and NAb structure are reported for Day 0 and Day 3 in Supplemental Tables 2 and 3, respectively. Most NAbs detected in survivors on Day 0 of ICU admission categorize as Class 3 (non-ACE2 blocking), while the most abundant NAbs in non-survivors belong to Class 4 (a different type of non-ACE2 blocking NAb). When assessing the number of counts that align to the non-ACE2 blocking C135 antibody, a SARS-CoV-2-specific NAb, survivors had significantly more (15,059.4) than non-survivors (1,412.7, p=0.016) on Day 0 but not on Day 3, suggesting an initial phase of NAb production may be critical for surviving COVID-19 ICU admission. All other distinct SARS-CoV-2 antibodies (as described in **Supplemental Table 1**) had no difference across survivors vs. non-survivors or across time points.

Top NAbs identified as unique to survivors and non-survivors on Day 0 are presented in **Table 2** (a full list is presented in **Supplemental Table 4** for survivors and **Supplemental Table 5** for non-survivors). Among survivors, most antibodies categorize as non-ACE2 blocking Class 3 while many of the top antibodies found in non-survivors were categorized as mainly Class 4 with some other classes of antibodies having fewer reads. In order to understand how non-ACE2 blocking NAbs would interact with the SARS-CoV-2 spike protein, we carried out MD simulations on the complex after modeling the CDR3 sequence most frequently found in survivors into an existing cryo-EM structure and report our findings as follows.

**Table 2.**
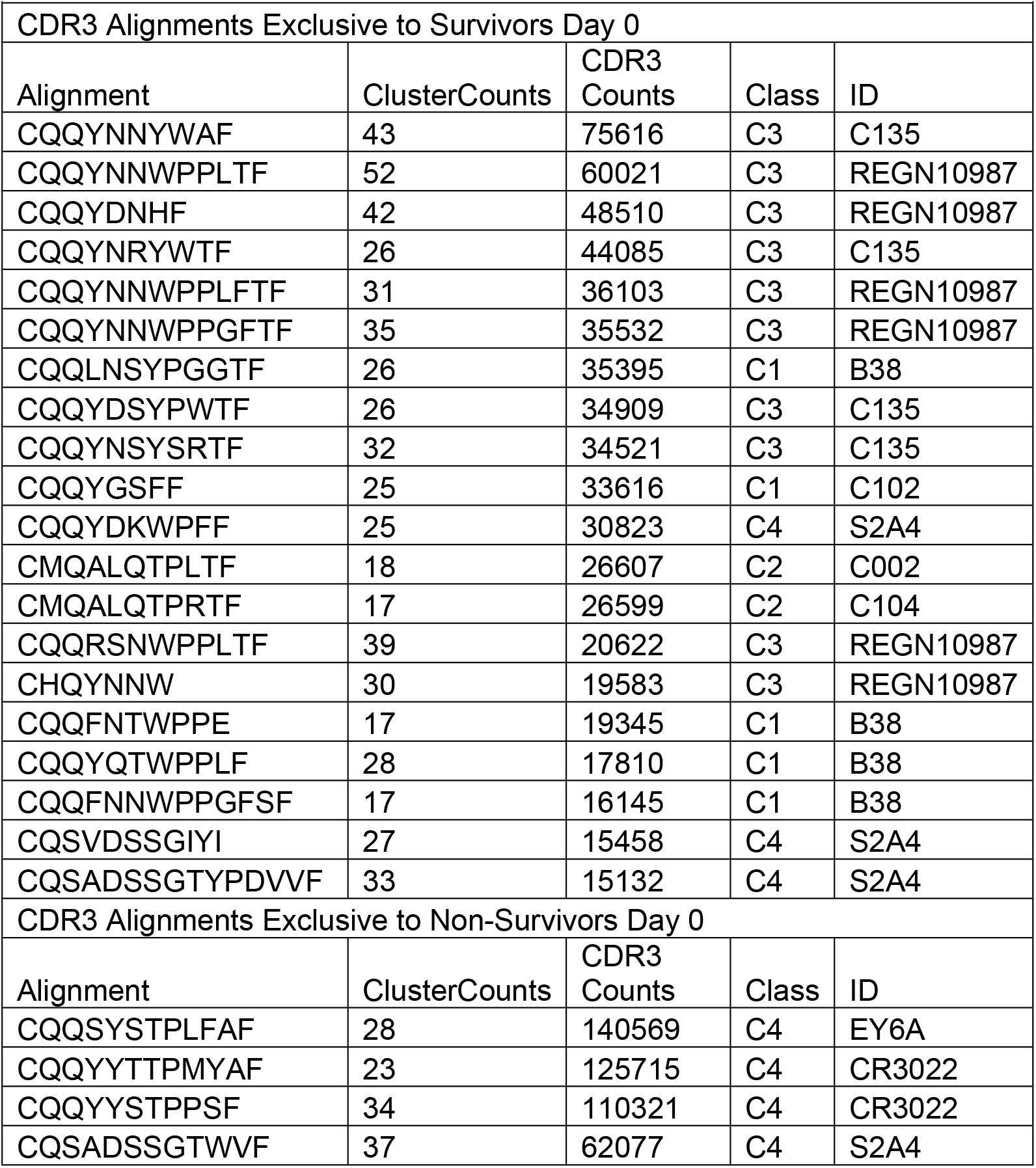
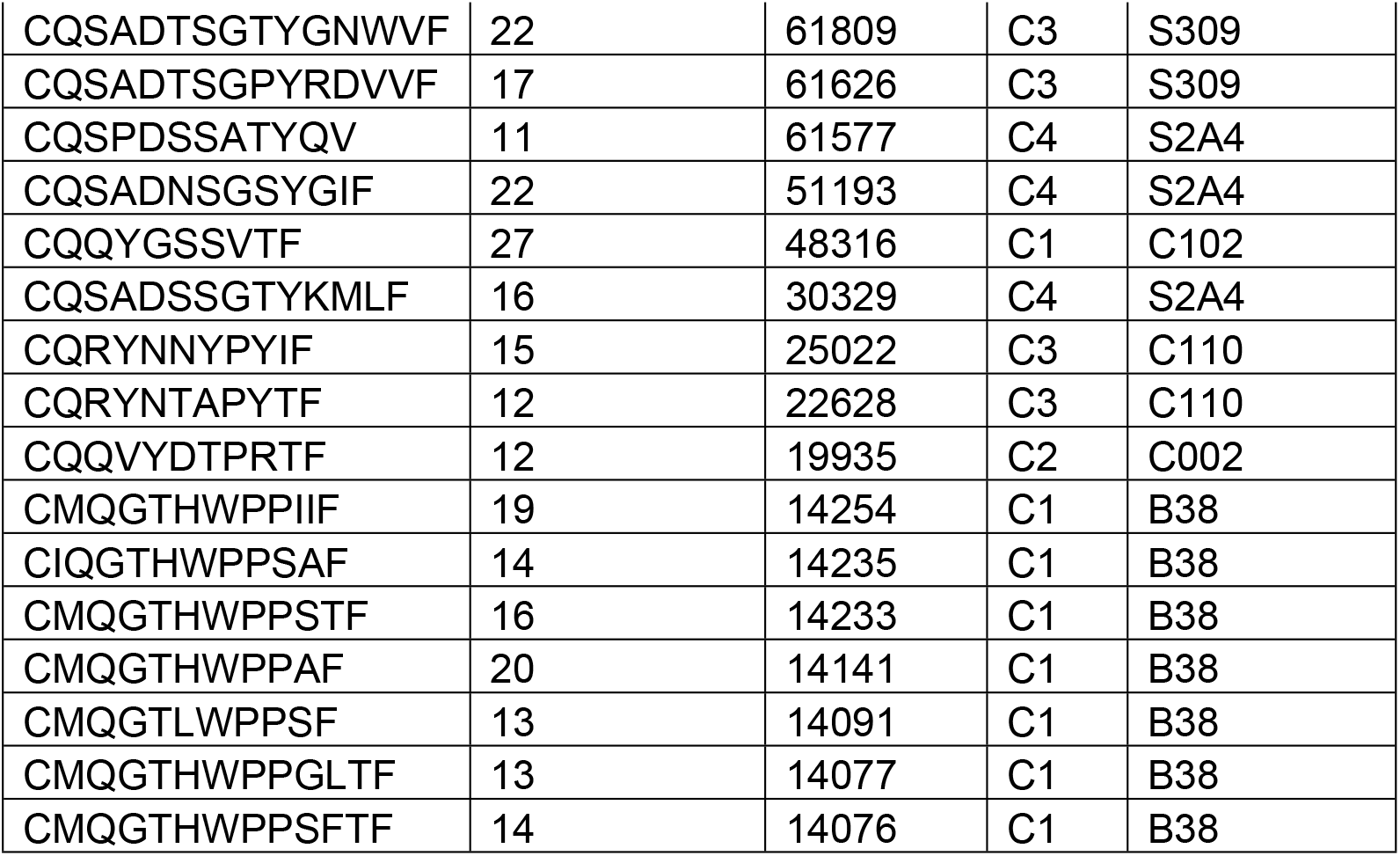
Top 20 CDR3 Alignments that were exclusive to survivors and non-survivors on ICU Day 0. The alignment is the common sequence to which a number of CDR3 from each patient aligned (ClusterCounts). The CDR3 counts represent the number of times a CDR3 mapped from the survivors or non-survivors. The class and ID are defined^22^ and references in Supplemental Table 1.

### Non-ACE2 blocking C135 NAbs can still preclude ACE2 binding

The cryo-EM structure reported for the spike/NAb C135 Fab fragment (PDB: 7k8z) shows that binding sites for the Fab fragment and the ACE2 receptor domain do not overlap.^22^ However, the 7k8z model built for C135 represents only about one half of its Fab fragment and about one-sixth of its entire NAb IgG1.^22^ The ACE2 receptor domain also represents only a fraction of the neutral amino acid transporter (N^0^AT1)/full-length ACE2 complex, which is a membrane-bound dimer that contains two ACE2 receptor domains.^35^ To determine whether binding of the intact NAb C135 IgG1 to the RBD of the spike trimer interferes with the binding of ACE2 receptors attached to the host cell membrane, we considered (1) the size of the intact NAb IgG1, and (2) the geometry and curvature of the host membrane and membrane-bound N^0^AT1/ACE2 dimer, as discussed previously.^17^

To represent the complete size of the NAb C135 IgG1, we used the monoclonal therapeutic anti-PD1 (Parkinson Disease) full-length IgG4 structure recently reported to represent the size of NAb C135 IgG1 in our modeling,^36^ although the elbow angle between the constant domain and Fab fragment may differ (**Fig. 1A**). We modeled one, two, or three complete NAbs C135 IgG1 bound to one, two, or three RBDs of the spike trimer in its two different conformations (**Fig. 1**). The first conformation was taken from the PDB: 6vxx coordinates, which represents the fully closed central pore with all three RBDs in the “down” position (**Fig. 1B, Video 1**).^37^ The second conformation is the symmetrized version of PDB: 6vyb (*i*.*e*. “6vyb”), which represents a partially opened central pore (open-1) with all three RBDs in the “up” position after rotating two “down” positioned RBDs (**Fig. 1C, Video 1**).^17,37^ Another fully opened conformation (open-2) was observed, which was essential for the productive host-viral membrane fusion reaction.^17,38^

The maximal dimension of the intact NAb C135 IgG is comparable with that of the ectodomain of the spike protein (**Fig. 1D, Video 1**). When one NAb C135 IgG binds one RBD of the fully closed spike trimer or to one “down” RBD in the mixed up-down asymmetric spike trimers, the large size of the NAb C135 IgG sterically blocks the virus from approaching the host membrane and thereby prevents the viral spike protein from binding to the N^0^AT1/ACE2 dimer (**Fig. 1D, Video 1**). Therefore, binding of a single NAb C135 IgG1 to one “down” positioned RBD (as well as multiple NAb C135 molecules) is sufficient to completely block binding of the N^0^AT1/ACE2 dimer and thereby prevents the ACE2-dependent host-viral membrane fusion (**Fig. 1D**).

### Non-ACE2 blocking NAb C135 can open the spike trimer and block membrane fusion

As long as one RBD is in a “down” position, the bound NAb C135 IgG1 will prevent the virus from approaching any host cell membrane from the direction of the occupied RBD. However, NAb C135 binds RBD independent of its conformational state (**Fig 1**), and when bound to an “up” positioned RBD, it will not prevent the virus from approaching the host cell membrane. If this RBD already has an ACE2 receptor bound, the Nab is still able to bind (**Fig. 2A**). Binding of either a non-ACE2 blocking NAb or the ACE2 receptor can shift the down-to-up equilibrium of the neighboring RBDs and thereby cooperatively open the central pore of the spike trimer (**Fig. 1, Video 1**). Therefore, this NAb can permanently disarm the triggering mechanism of the spike trimer with or without assistance of the ACE2 receptor. As MD simulations described below, the binding of NAb C135 IgG1 weakens inter-subunit interactions that constrain the RBDs in “down” positions within the spike trimer.

**Figure 2.**
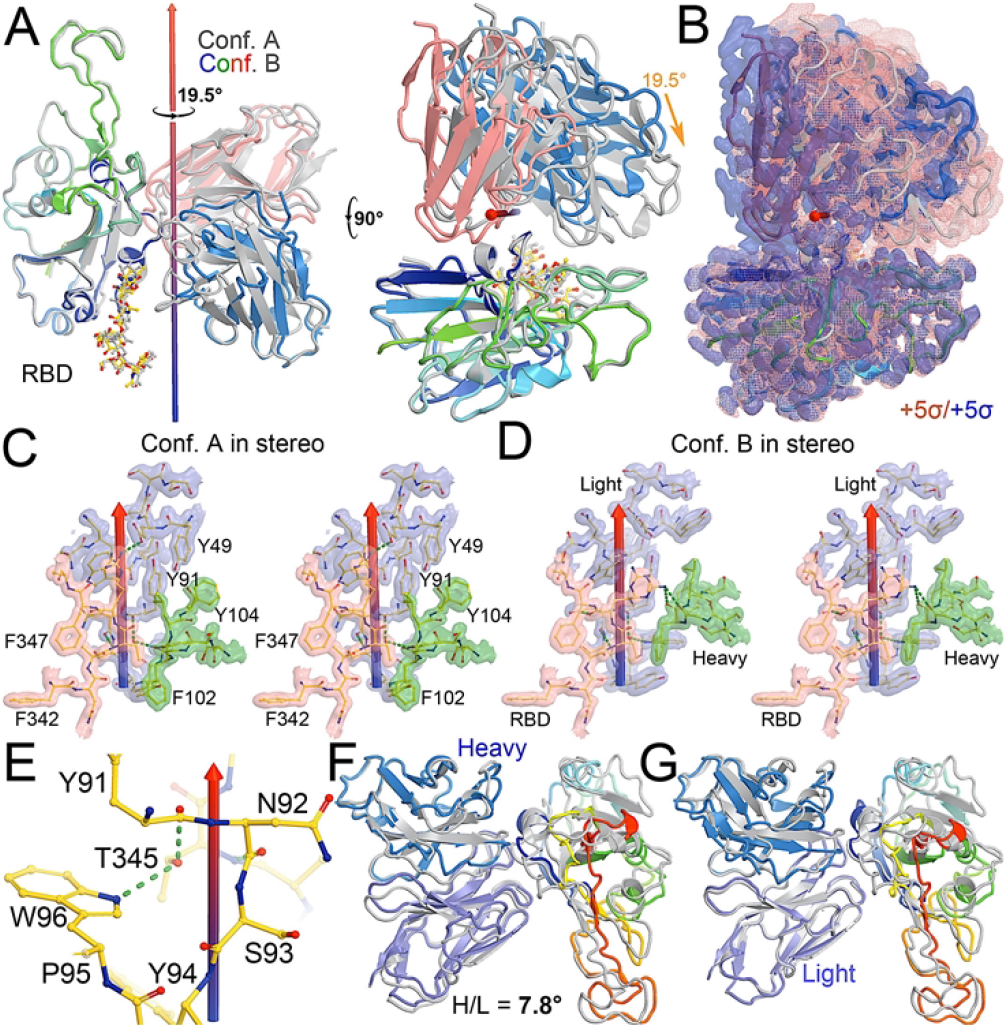
Comparison between conformations A and B of the NAb-spike complex. (A) Two orthogonal views of comparison with rotational axis and angle indicated upon alignment of the RBD (rainbow color) between conformation A (grey) and B (multicolor). (B) Corresponding MD-ESP maps contoured at +5σ. (C) Interactions of the RBD and heavy/light chains near the rotational axis in stereodiagram of conformation A. (D) Conformation B in stereodiagram. (E) Conserved interaction surrounding T345 in the two conformations. (F) Alignment of heavy chain between the two conformations. (G) Alignment of light chain.

The fusion reaction of two negatively charged membranes is an energetically unfavorable process that is made possible when coupled with the highly exergonic conversion of the prefusion to postfusion states of the S2 fragment of the spike protein. A single spike trimer can fuse the two membranes but cannot open a connecting pore between them to transfer genetic materials from the virus to the host cell.^17^ This requires formation of supercomplexes containing 3 spike trimers and 6 N^0^AT1/ACE2 dimers, involving binding of two NAbs to each ACE2 dimer.^17^ Modeling of the full-length NAb C135 IgG1 and full-length N^0^AT1/ACE2 dimer shows that two IgG1 molecules have steric clashes that enable the NAb to block this step of membrane fusion process (**Fig. 2, Video 2**).

### Two stable states of the Nab C135-RBD complex are revealed by MD simulations

After a three-step minimization of the NAb C135-RBD complex, MD simulations were carried out for 465 ns and coordinates were written every 200 ps. During the MD simulations, the complex rapidly evolved away from the starting configuration (conformation S) via short-lived intermediates and rapidly settled into two stable configurations (conformation A and B) according to a root-mean-square fluctuation (rmsF) analysis in reference to either frame 1 or frame 500 (or to frame 1000, data not shown). Conformation A begins at frame 360 and lasts for 98 ns. Conformation B begins at frame 849 and lasts for 149 ns (**Fig. 1A**). In our initial analysis, we combined the two conformations into a single state that represents 52% of the total trajectory frames. Each of the two conformations in the MD trajectories could have persisted for much longer, but we were unable to reliably estimate their duration. However, during interpretation of the MD-derived electrostatic potential (ESP) maps, we found two clear and closely related conformations (**Fig. 3, 4, S5**).

**Figure 3.**
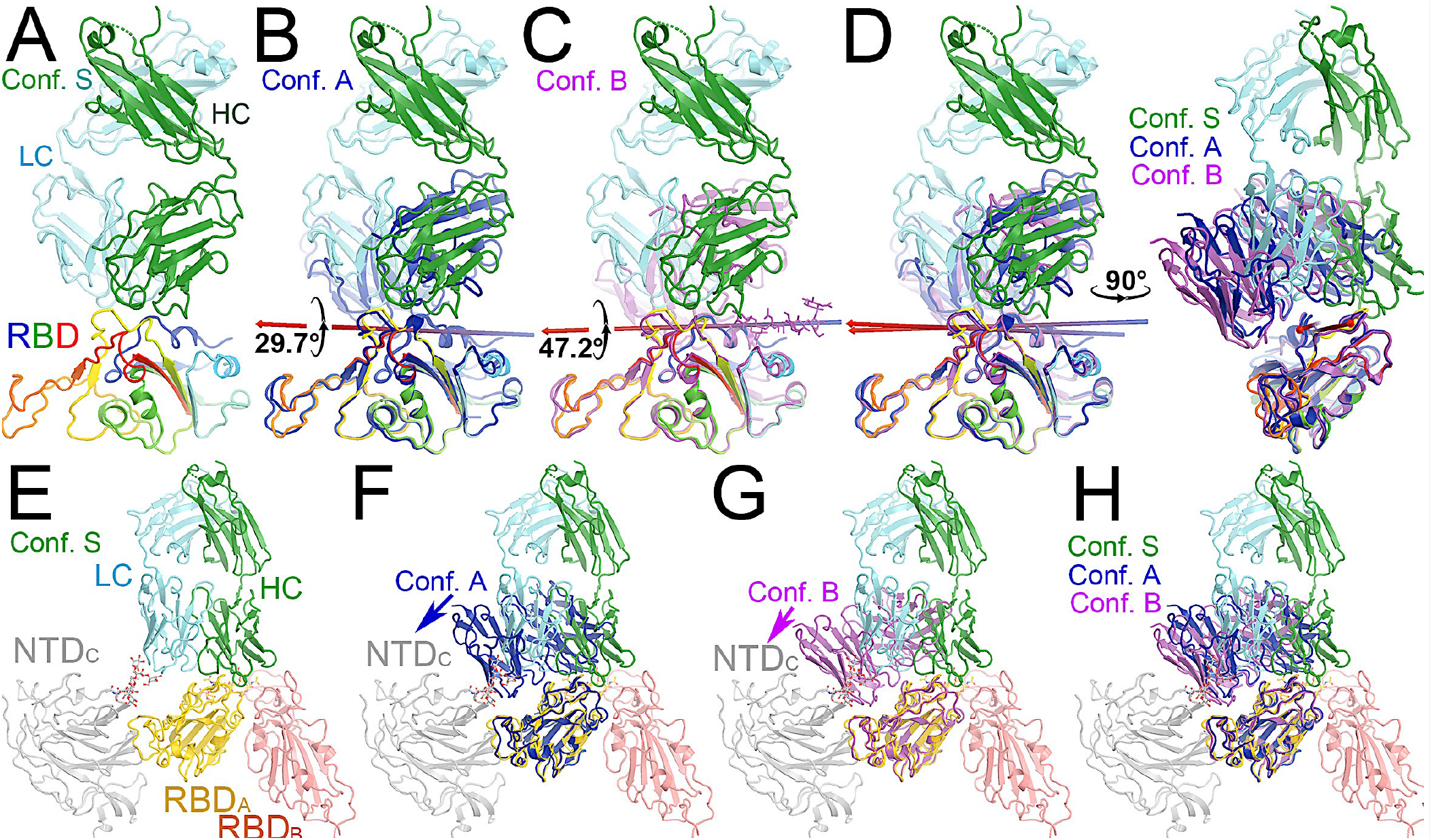
Comparison of MD-derived conformation A (blue) and B (magenta) with the starting conformation S (multicolor). (A) Starting conformation S. (B) Superposition between conformations S and A. (C) Superposition between conformations S and B. (D) Two orthogonal views of conformations S, A and B. (E) Conformation S in context of the spike trimer with NTD and RBD of neighboring subunits included. (F) Superposition of conformations S and A in the context of the spike protein. (G) Superposition of conformations S and B in the trimer. (H) Compiled view of conformation S and A/B in the trimer.

**Figure 4.**
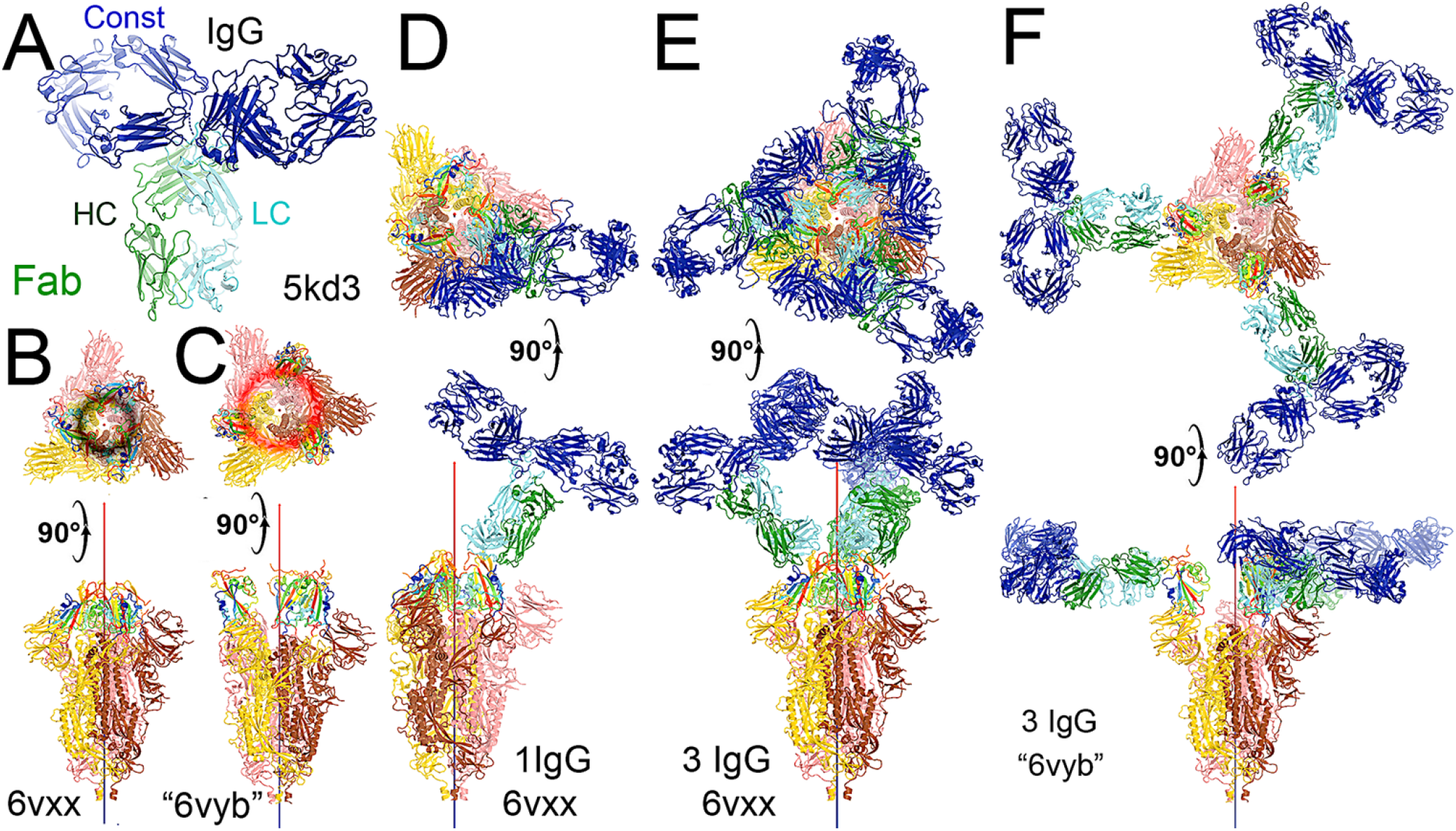
Binding of the full-length NAb C135 to the spike trimer in two conformations. (A) Using the full-length anti-PD1 IgG4 to represent the full-length C135, constant domains are in blue, heavy chain in green, and light chain in cyan. (B) Two orthogonal views of the closed spike trimer (from PDB: 6vxx coordinates). (C) Two orthogonal views of the fully open (open-1) spike trimer (using symmetrized PDB: 6vyb coordinates). (D) Binding of one NAb C135 to the closed spike trimer in two orthogonal views. € Binding of three NAb C135 to the closed spike trimer in two orthogonal views. (F) Binding of three NAb C135 to the open spike trimer.

When MD-derived ESP maps are visualized in continuous contouring levels of 10σ, 5σ, 2.5σ, and 1.25σ, the NAb C135/RBD interface remains the most ordered part of the complex (**Fig. S5**) This region of the equilibrated structure has the smallest rmsF and the smallest atomic B-factors after the fitted models were refined. While the interface area buried between the RBD and NAb C135 is 8,132 Å^2^ (heavy chain, 4,536 Å^2^, light chain, 3,596 Å^2^, including 2,387 Å^2^ between the heavy chain and N343 glycans) for conformation A, and 6,476 Å^2^ (heavy chain, 1681 Å^2^, and light chain 4794 Å^2^, including only 432 Å^2^ between the heavy atom and N343 glycans) for conformation B, multiple hydrogen (H)-bonds are observed. Inter-subunit H-bonds include R346 of the RBD and E50 of light chain (numbering of antibody residues in this study is according to their genomic coding sequence, but it does not include gaps and insertions in the aligned antibody structure) and T346 of the RBD with both the W96 sidechain and Y91 backbone carbonyl and N92 backbone carbonyl of the light chain (**Fig. 3E**). There are also extensive interactions between the heavy chain and the glycan units attached to N343 of the RBD when the MD-derived ESP maps are viewed at reduced contouring levels (**Fig. 3G**). In conformation B, the H-bond pattern of T346 of the RBD remains unchanged relative to conformation A, but R346 of the RBD is repositioned to form H-bonds in one of the two sidechain rotamers with the carbonyl groups of both G101 and F102 of the heavy chain (**Fig. 4C, 4D**).

The two conformations differ by a rotation of 19.5° around an axis passing through the conserved H-bond network of T346 (**Fig. 4A**). This rotation is evident in the two MD-derived maps even before the maps were interpreted and the coordinates were built (**Fig. 4B**). In addition, there is a small rigid rotation of the heavy and light chains within NAb C135 (**Fig. 4F, 4G**), revealing that dynamics of this complex are largely associated with rigid-body rotations of domains with a small rmsF value of ∼1.8 Å for the entire complex in MD trajectories of each conformational state (**Fig. 4**).

### Functional roles of glycans in binding of NAb C135 to the RBD

Viruses often hijack host glycosylation machinery to hide epitopes of their spike proteins. The majority of NAbs do not recognize glycans because these are components of host cellular proteins and specified by the individual’s blood type. We found, unexpectedly, that R56, N57, and Y59 of the heavy chain of NAb C135 make favorable transient interactions with glycosylated residue N343 for a significant duration of MD trajectories only in conformation A (**Fig. 3G**). Likewise, the light chain of NAb C135 would also interact with glycosylated N165 of the neighboring N-terminal domain (NTD) (**Fig. S3, S6**), which was not included in our current MD simulations. In fact, interactions of a given NAb with glycans may help to recruit the NAb before it fully recognizes the underlying epitope, *i*.*e*. this binding represents the first step of antibody-epitope recognition. Much of the heavy chain-glycan interactions are lost in the most persistent, and likely more stable, conformation B. Our MD simulations reveal stable antibody-epitope interactions after Nab C135 peels away from the N343 glycans and likely also the N165 glycans according to the observed domain rotations.

Comparison with the starting configuration (conformation S) taken from the NAb C135/spike complex structure reveals how NAb C135 fully recognizes the epitope of the RBD. Recognition involves a rotation of NAb C135 in conformation A by 29.7° and then conformation B by 47.2° with two closely related axes that are similar to the rotational axis separating the conformations (**Fig. 5, Video 3**). The direction of these NAb C135 rotations relative to the RBD would place the light chain of the NAb in a partially overlapping position with both its NTD and N165 glycans within the NAb C135/spike complex. Because the RBD interacts with the N165 glycans of a neighboring NTD within the complex (**Fig. S3, S6**), an initial binding of NAb C135 weakens this interaction and gradually pushes the NTD away from the spike RBD to facilitate full NAb C135-epitope recognition. This frees the RBD from constraints of its inter-subunit interactions within its trimer and allows the down-to-up conversion of the RBD.

**Figure 5.**
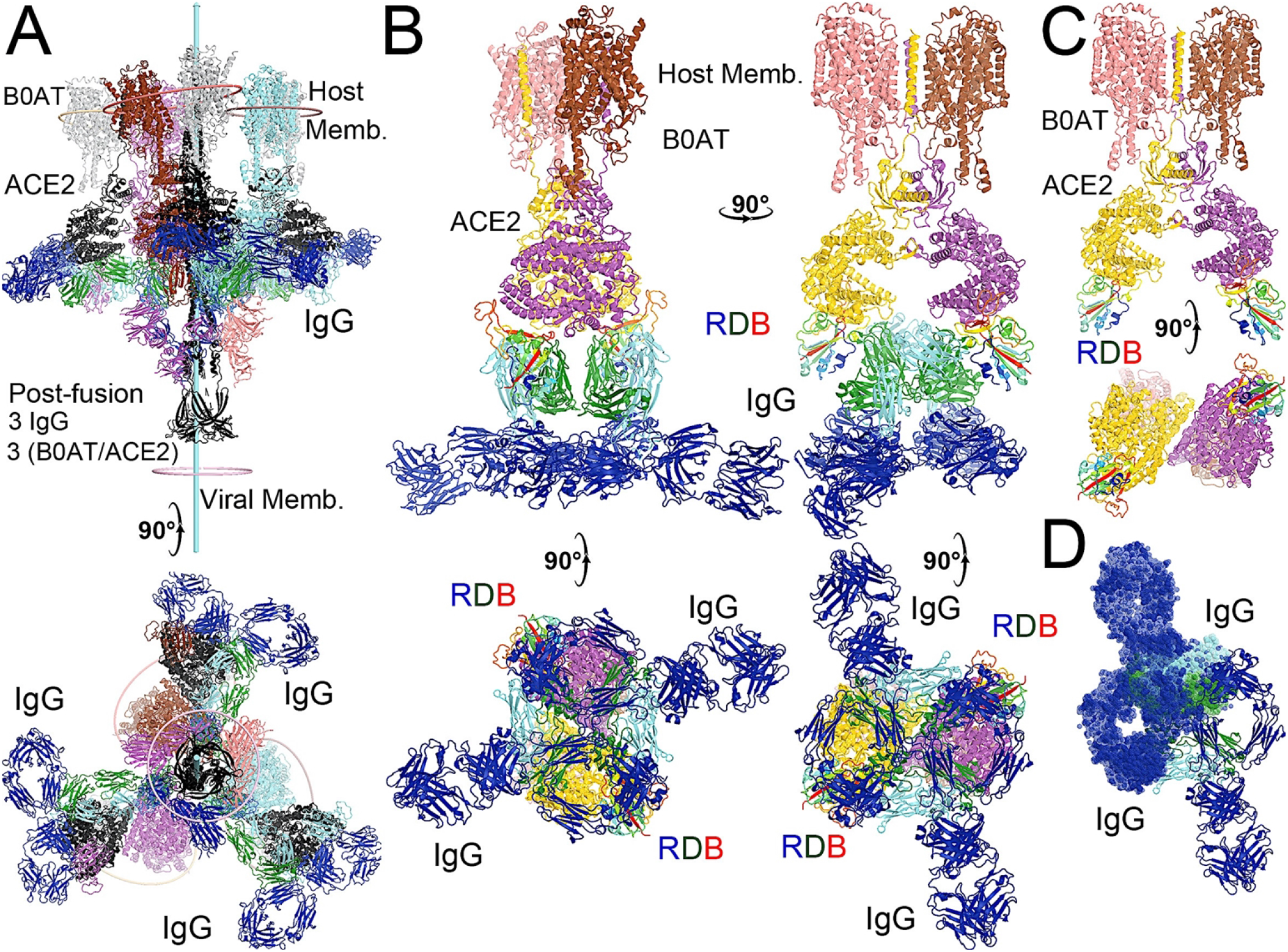
Modeling of the full-length NAb C135 with the full-length N0AT1/ACE2/spiker supercomplexes. (A) Two orthogonal views of 3 NAb C135 (blue, forest, and cyan) and 3 B0AT1/ACE2 (magenta, pink, and grey) dimers bound to one open spike trimer (black for the extended S2 central stalk, and multicolor for NTD and rainbow colors for RBDs). (B) Two NAb C135 bound to the B0AT1/ACE2 dimer in the opened ACE2 cleft in four orthogonal views. (C) B0AT1/ACE2 dimer with the spike RBD (rainbow color), but without the model NAb C135. (D) Two simultaneously modeled NAb C135 on one ACE2 dimer showing steric clashes between the constant domains and heavy/light chains of NAb C135.

The transition from initial binding of the NAb C135 to the spike trimer to full recognition revealed in our MD simulations involves the gradual engagement of the NAb light chain via specific hydrogen bonds and extensive hydrophobic interactions, as the heavy chain become less engaged. The interface between the heavy chain of the NAb and the spike RBD includes an induced-fit recognition of the NAb in variable loop 3 of the heavy chain, *i*.*e*. the conformation of this loop in the complex differs significantly in the unbound NAb (**Fig. S6**).

The NAb C135/spike complex obtained through our MD simulations only partially represents the full NAb/epitope recognition mode because formation of the stable RBD-NAb C135 complex in its full recognition mode provides greater rotational freedom of the spike RBD relative to the remaining part of the spike trimer. With this increased rotational freedom, the stable complex gradually becomes invisible in the cryo-EM map, indicating significantly reduced local resolution. In fact, the “up” positioned RBDs always have relatively low local resolution with respect to the more stable “down” positioned RBDs in all cryo-EM structures of the spike trimer and its complexes with NAbs or ACE2.

## Discussion

Deep RNA sequencing is a novel tool to identify antibodies elicited in response to infections, including COVID-19. Typical identification of antibodies is time consuming.^39^ In addition, antibodies identified from plasma do not provide as much detail as is gleaned from the RNA sequencing data among Ig classes and the actual amino acid sequence of the antibody. By identifying antibodies that are unique to COVID-19 survivors, we propose the CDR3 sequences most likely to positively impact infection. Utilizing this amino acid sequence created *in vivo* by a critically ill patient for models of a NAb/spike complex, we are able to establish the molecular details of the binding interaction via MD simulations.

The majority of prior work assessing immunoglobulin light chain types of lambda and kappa has been limited to hematologic malignancies and HIV infection.^40-42^ The impact of an increase of Ig lambda at ICU Day 3 in survivors is unknown. However, this could also be used as a potential marker of recovery. In addition, the use of an RNAseq-based workflow could enhance the study of class types in antibodies not only during infectious disease but also to bolster previous findings in hematologic malignancies and transplant medicine.

Classes of NAbs (Class 1, 2, 3, 4) against SARS-CoV-2 are catalogued based on their interactions with the spike protein,^22^ especially related to the conformation of the spike RBD to which they bind - (1) NAbs encoded by VH3-53 gene with short CDRH3 loops and bind only to “up” RBDs to block interaction with the ACE2 receptor; (2) NAbs that bind both “up” and “down” RBDs to block ACE2 and contact RBDs from adjacent spike subunits; (3) NAbs that bind outside the ACE2 site and recognize both “up” and “down” RBDs; and (4) NAbs that bind only “up” RBDs but do not block ACE2 binding. When assessing top counts of unique NAbs on Day 0 between COVID-19 survivors and non-survivors, the most abundant NAbs among survivors belong to Class 3 (including the non-ACE2 blocking C135-type described above), while the most abundant NAbs among non-survivors belong to both Class 2 and Class 4. Unexpectedly, there is no detectable difference in Class 1 ACE2-blocking NAb production between surviving and non-surviving patients. Class 4 NAbs bind the opposite side of the RBD that does not overlap with either Class 1 or Class 3 NAbs, requiring “up” positioned RBDs to expose the otherwise buried epitopes. According to our atomistic modeling, the binding of this class of NAbs could actually sterically block the partially open central pore of the RBDs and thereby physically obstructing the extension of the S2 central stalk during the prefusion-to-postfusion transition. However, this Class 4 NAbs are not expected to prevent the virus from approaching the host cell membranes, unlike Class 3.

Antibodies that bind at the ACE2 site on the spike RBD are also known to be influenced by viral mutations.^4^ With the proliferation of variants^43^ as the pandemic continues, it is important to ensure that antibodies exist that are not disrupted by mutations at the ACE2-binding motif. Class 3 and 4 antibodies bind to an epitope orthogonal to the ACE2 binding site and are known to attenuate the virus in cells that overexpress ACE2.^4^ ACE inhibitors are known to increase ACE2 expression^44^ but there has been no definitive correlation with patient outcome.^45^ This lack of impact of ACE inhibitors may be due to a different class of antibodies that a patient produces in response to infection. Unfortunately, this sample size is not sufficient to assess the impact of pre-infection ACE inhibitor use. However, since NAb C135 binds to the spike protein distal to the ACE2 binding site and is produced in substantially higher levels in survivors, this suggests a unique ‘ACE2-independent’ mechanism of antibody mediated protection.

The spike protein forms a trimer where all must be in the “up” position for RBD binding to ACE2.^46^ Simulations of spike trimers showed its down-to-up transition as well as locked, stabilized “down” states, where acidic-pH confers stability to the locked structure.^46^ Class 1 and 4 NAbs are only able to bind in the “up” conformation, while Class 2 and 3 NAbs are able to bind the spike protein in both the “up” and “down” conformations, which is the most effective for neutralizing the virus.^9,10^ It is also proposed that utilizing a spike protein in the “down” position may result more effective vaccines.^47^

Cellular transmission of both SARS-CoV and SARS-CoV-2 are pH (*i*.*e*. protonation state) dependent, as is that of many other coronaviruses.^48,49^ At low pH, viruses can transmit locally by cell-to-cell spreading through cell fusion and endocytosis.^50,51^ However, different coronaviruses have distinct pH sensors that exhibit unique pH dependencies. For example, helical repeat 1 (HR1) and helical repeat 2 (HR2) of the central stalk of the influenzae spike protein (i.e., hemagglutinin) become extended as pH decreases,^52^ which is caused by changes in protonation states of its pH-sensing residues.^51,53,54^ However, lower pH values stabilize the spike protein of SARS-CoV-2 in the fully closed conformation with its three RBDs in the “down” position and its HR1 and HR2 of the S2 central stalk in a non-extended conformation.^55^

Patients that are critically ill are typically acidotic. Thus, NAbs such as the Class 3 C135 NAb, which are able to efficiently bind the “down”-positioned spike RBD and to remain bound should a pH-driven conformational change occur, could be ideal therapeutics for ICU patients. The down-to-up pH-dependent equilibrium of the spike protein is also highly sensitive to spontaneous mutations. For example, the spike protein of the Omicron variant is in an open conformation at pH 7.5 (100% with one RBD “up” and two RBDs “down”) whereas the spike protein of the prototype SARS-CoV-2 (Wuhan strain plus the D614G mutation) is 50.3% in the open conformation and 49.7% in the fully closed conformation,^56^ highlighting different mechanisms of cellular transmission between the two variants. Moreover, mature virions of SARS-CoV-2 are released from acidified Golgi via exocytosis during which the spike protein is also stabilized in the closed state.^57,58^ Interestingly, fatalities from severe infection often result from non-local cell-to-cell transmission that is dependent on ACE2 while initial SARS-CoV-2 infection involves mainly local cell-to-cell transmission that is largely independent of ACE2.^59^ Thus, ACE2 blocking NAbs likely have little effect on this initial step of local cellular transmission, where only non-ACE2 blocking NAbs can exhibit dual functions, inhibiting both the local and non-local cellular transmission upon their binding. This highlights the importance of including non-ACE2 blocking NAbs in therapeutic treatments of SARS-CoV-2. Moreover, NAbs elicited in response to an immunogen of the closed state-stabilized spike trimer are distinct from those produced in response to the mRNA-based or protein-based RBD-only proteins used in current vaccine designs.^60^ Our new understanding the molecular mechanism(s) of non-ACE2 blocking NAbs could help to design the next generation of immunogens for more effective vaccines.

Some engineered antibodies appear to already utilize sites that mimic a cocktail of both Class 1 and 4 antibodies.^61^ However, based on our current study, it appears that focusing on Class 3 antibodies, with further structural work, could reveal the ideal COVID-19 antibody. The widely reported Regeneron Antibody^7^ is one of Class 3, but distinct from C135,^22^ which was also elevated in COVID-19 survivors relative to non-survivors on ICU Day 0. In one patient with a low level of C135 compared to other survivors, an increase in another Class 3 antibody similar to REGN10987 was found (**Supplemental Tables 2 and 3**), highlighting the importance of Class 3 NAbs. This suggests that if patients with low levels of Class 3 antibodies, specifically C135, are identified by RNAseq on ICU Day 0, they could be treated with other commercially available Class 3 NAbs. With further analysis of antibody profiles from patients with new SARS-CoV-2 variants and extensive structural assessments, more versatile monoclonal antibodies could be produced using recombinant techniques for treatment of critically ill patients.

Such a need is illustrated when considering a SARS-CoV-2 variant such as Omicron, with 29 single amino acid mutations in its spike protein, as well as several insertions/deletions. Nearly half of these mutations occur within the spike RBD and several at the RBM, which could alter interactions between the RBD and ACE2 that are essential for viral reentry to the host cells. Since prior biophysical studies have suggested that conformational flexibility of the RBM is essential for the stability of the RBDs in the “down” state, Omicron mutations could destabilize the “down” state, attenuating the ability of NAbs to promote this conformation and facilitating viral entry into cells. RNAseq from patients who survive an infection of future SARS-CoV-2 variants could elucidate antibodies created to solve this problem, particularly if those NAbs are capable of binding to both “up” and “down” spike conformations (*i*.*e*. such as C135). This scenario takes on heightened importance as a significant body of work has shown that NAbs specific for “wild-type” SARS-CoV-2 are less effective against variants. Mutant spike protein from SARS-CoV-2 is resistant to sera from convalescent or mRNA vaccine recipients but not to patients infected and then vaccinated.^62^ In addition, SARS-CoV-2 variants have shown some NAbs to be less effective, requiring additional vaccine doses and highlighting the importance of both antibody development and vaccination in the care of COVID-19. RNAseq with analysis for CDR3 sequences could, if optimized, uncover novel antibodies in real-time as patients are hospitalized with critical illness from future variants. RNAseq could also assess response to vaccines, especially in immunosuppressed patients,^63^ providing insight into a cocktail of antibodies that may be beneficial in treating COVID-19, in addition to ubiquitous vaccination.

## Declarations

### Funding

This study was supported by funding from the US National Institutes of Health: T32 HL134625 (AMF), R35 GM118097 (AA), P20 GM103652 (WGF, AA, SFM), R01 GM 127472 (WGF), R35 GM124911 (CTL), P20 GM121344 (GJN), R01 GM 136815 (VSB, GPL), R35 GM 142638 (SFM).

### Conflicts of Interest

Some of the techniques described here have been submitted in provisional and utility patent applications. Early results from this data were presented at the virtual Shock Society Meeting 2021. Sean Monaghan is founder of company Alcini, LLC.

### Ethics approval

Institutional Review Board Approval # 411616

### Availability of data

See online supplement for publication of extensive data.

### Code availability

All code used is cited in the text and custom scripts are available if requested

### Author Contributions

Drs. Fredericks and Monaghan had full access to all of the data in the study and take responsibility for the integrity of the data and the accuracy of the data analysis.

*Concept and design:* Fredericks, East, Lisi, Monaghan

*Acquisition, analysis, or interpretation of data:* Fredericks, East, Wang, Liu, Shi, Maschietto, Cohen, Lefort, Nau, Levy, Liu, Wang, Maschietto, Batista, Lisi, Monaghan

*Drafting of the manuscript:* Monaghan, Fredericks.

*Critical revision of the manuscript for important intellectual content:* Fredericks, East, Wang, Liu, Shi, Maschietto, Ayala, Cioffi, Cohen, Fairbrother, Lefort, Nau, Levy, Lisi, Monaghan

*Statistical analysis:* Fredericks, East, Wang, Fairbrother, Lisi, Monaghan

*Obtained funding:* Fredericks, East, Ayala, Fairbrother, Lefort, Nau, Batista, Lisi, Monaghan

*Administrative, technical, or material support:* Ayala, Cioffi, Fairbrother, Lefort, Nau, Levy, Lisi, Monaghan

*Supervision:* Ayala, Fairbrother, Levy, Batista, Lisi, Monaghan

